# Versican binds collagen via its G3 domain and regulates the organization and mechanics of collagenous matrices

**DOI:** 10.1101/2022.03.27.485990

**Authors:** Dongning Chen, Yu Du, Jessica Llewellyn, Arkadiusz Bonna, Biao Zuo, Paul A. Janmey, Richard W. Farndale, Rebecca G. Wells

## Abstract

Type I collagen is the most abundant structural protein in the body and, with other fibrillar collagens, forms the fibrous network of the extracellular matrix (ECM). Another group of ECM polymers, the glycosaminoglycans (GAGs) and GAG-modified proteoglycans, play important roles in regulating collagen behaviors and contribute to the compositional, structural and mechanical complexity of the ECM. While the binding between collagen and small leucine-rich proteoglycans (SLRPs) has been studied in detail, the interactions between collagen and the large bottlebrush proteoglycan versican are not well understood. Here, we report that versican binds collagen directly and regulates collagen structure and mechanics. Versican colocalizes with collagen fibers *in vivo* and binds to collagen via its C-terminal G3 domain (a non-GAG-modified domain present in all known versican isoforms) *in vitro*; it promotes the deposition of a highly-aligned collagen-rich matrix by fibroblasts. Versican also shows an unexpected effect on the rheology of collagen gels *in vitro*, causing decreased stiffness and attenuated shear strain stiffening, and the cleavage of versican in liver results in reduced tissue compression stiffening. Thus, versican is an important collagen binding partner, playing a role in modulating collagen organization and mechanics.

## Introduction

Type I collagen, the most abundant protein in the body, forms a dynamic fibrous network that plays a critical role in maintaining normal cell and tissue structure, function, and mechanics. The structure and mechanics of the collagen fibrous network are interrelated [1] and are highly regulated by factors including cell-generated force [2] and other extracellular matrix (ECM) components such as fibronectin [3], glycosaminoglycans (GAGs), and proteoglycans (proteins modified by GAGs) [4][5][6]. GAGs are highly negatively charged and contribute to collagen mechanics by attracting water, thereby swelling and resisting compression [7]. GAGs and proteoglycans are important regulators of collagen-related fibroproliferative diseases such as inflammation, fibrosis and cancer [8][9] and understanding the relationships between these matrix components and collagen is thus important to understanding pathophysiology.

There are two families of matrix (interstitial) proteoglycans: small leucine-rich proteoglycans (SLRPs) and hyalectans (also known as large chondroitin sulfate proteoglycans). SLRPs, which include decorin, lumican, and fibromodulin, have core proteins of about 50 kDa with 1-4 GAG side chains. They have been well investigated as regulators of collagen fibrillogenesis and fiber organization; *in vivo* studies of tendon development have demonstrated a crucial role for SLRPs in regulating fiber size, morphology and organization [10]. Crystal structures for some SLRPs (showing horseshoe-like structures) and the use of solid-phase binding assays have led to the identification and modeling of binding sites between collagen and SLRPs [11][12]. The use of synthetic peptides containing different leucine-rich repeats (LRRs) showed that fibromodulin and lumican bind collagen via the same LRR [13]. The development of Collagen Toolkits (libraries of synthetic collagen mimetic peptides) has been a major step forward in mapping protein binding sites on collagen [14][15]. Taking advantage of this tool, fibromodulin was found to interact with the collagen MMP cleavage site as well as the KGHR sequence, which is involved with helical crosslinking [16].

Interactions between collagen and members of the hyalectan family are less well defined. Hyalectans are large proteoglycans with mainly chondroitin sulfate (CS) side chains that bind hyaluronic acid (HA). This group includes versican (~360 kDa core protein with multiple isoforms (Figure 1A) and 0-23 CS chains, depending on the isoform) and aggrecan (~250 kDa core protein with approximately 30 keratan sulfate and 100 CS chains) [17]. Aggrecan, found primarily in cartilage, interacts with both type I and II collagen via its keratan sulfate domain in a manner dependent on ionic strength, as defined using solid-phase binding assays [18]. Solidphase binding assays have also shown that versican can bind collagen [19], but the binding site has not been identified. Versican, unlike aggrecan, is found throughout the body. It has been studied as a modulator of diseases including hepatic and pulmonary fibrosis and thus its interactions with collagen are particularly important to understand [20][21]. We have recently highlighted differences in the effects of versican compared to other matrix proteoglycans on collagen behaviors; specifically, we showed *in vitro* that versican upregulates collagen fibrillogenesis and the alignment and contraction of engineered collagenous tissues, and that the presence of versican results in collagen matrices with fibers fused into large bundles [5].

**Figure 1.**
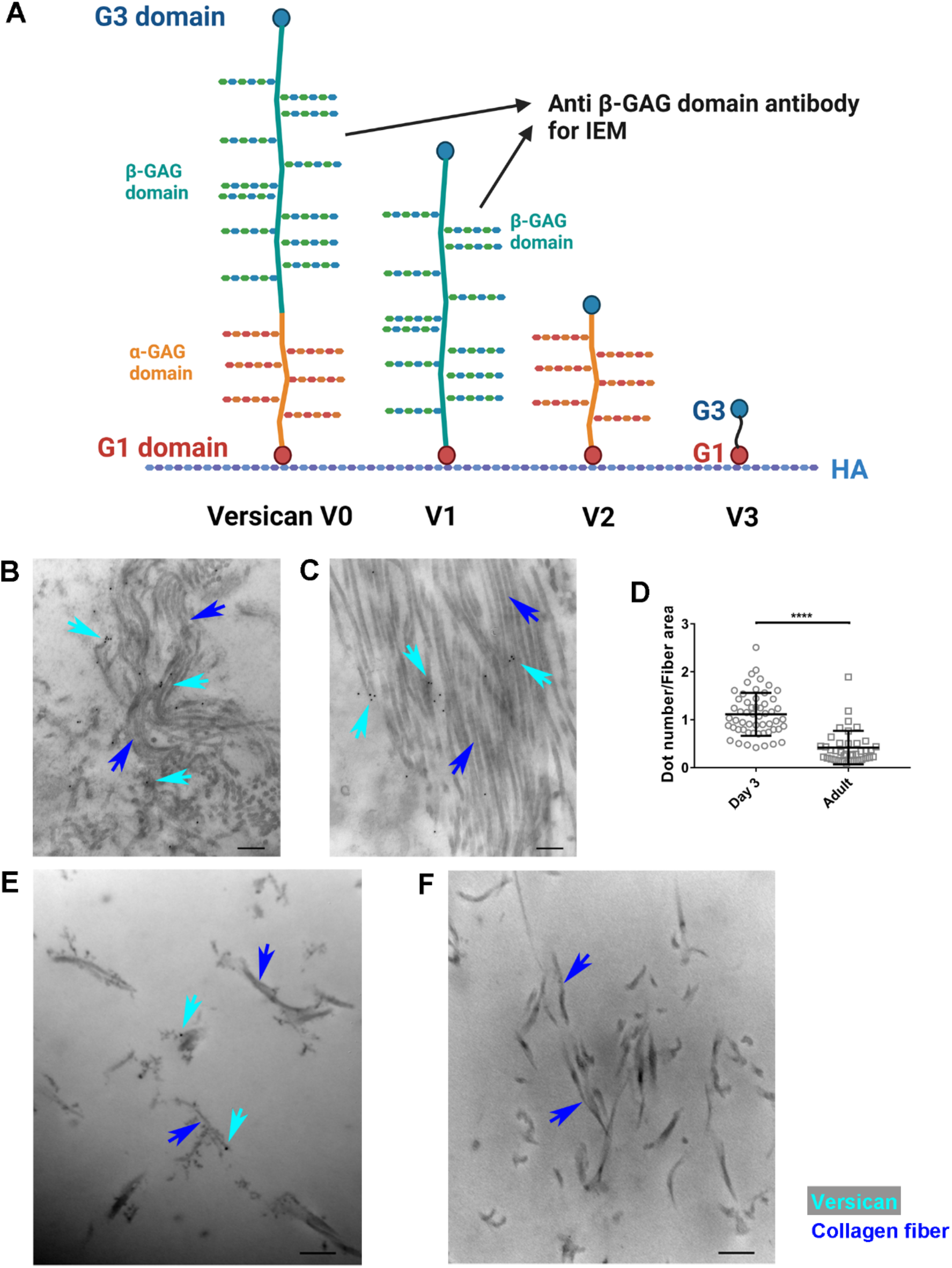
Versican co-localizes with collagen fibers in mouse extrahepatic bile duct and collagen gels. **A)** Schematic showing the different isoforms of versican and the location of the antibody used for IEM within the β-GAG domain. **B, C)** Representative IEM images showing localization of versican (black dots, indicated by teal arrows) on collagen fibers (blue arrows) in postnatal day 3 (B) and adult (C) mouse bile ducts. **D)** Quantification of the number of gold particles, indicating versican, per total fiber area. **E, F)** Representative IEM images showing the localization of versican in plugs of co-gelled collagen (blue arrows) and versican (black dots indicated by teal arrows) (E) or collagen alone (F). Data represent mean ± SD and were analyzed by unpaired t test, ****P<0.0001.

We report here the results of further investigation into the interaction between versican and collagen. We used solid-phase binding assays including the Collagen Toolkit to identify binding sites on both proteins. We also used fibroblast-derived matrices, versican-collagen co-gels, and rat liver tissue to demonstrate that these interactions are functionally relevant.

## Results

### Versican co-localizes with collagen fibers *in vivo*

We first examined the colocalization of collagen and versican *in vivo*. Versican and collagen are both found in the submucosa of the mouse extrahepatic bile duct during development and in the adult [22] and we therefore used immunoelectron microscopy (IEM) to determine whether they co-localize in this physiologic setting. The antibody used to detect versican recognizes an epitope in the β-GAG domain, such that it detects the large V0 and V1 isoforms (Figure 1A). In almost all cases, versican, which is indicated by a black dot (12 nm) on IEM images (Figure 1B, C), was closely apposed to collagen fibers. The versican signal decreased during bile duct development, as shown previously by immunostaining [22], and the the co-localization between collagen and versican similarly decreased (Figure 1D). IEM of collagen-versican gels confirmed the co-localization between versican with collagen fibers (Figure 1E, F). Thus, versican and collagen fibers co-localized both *in vitro* and *in vivo*.

### Versican core protein binds collagen via its G3 domain

To better define the interaction between collagen and versican, we first attempted to use the Biacore surface plasmon resonance system but found unacceptably high non-specific binding of versican to the control chip. We thus moved to traditional solid-phase binding assays. Plates were coated with native versican (referred to here simply as versican), native versican treated with chondroitinase ABC (ChABC) to remove GAGs (referred to here as core protein), recombinant versican V3 isoform (which contains only the N-terminal G1 and C-terminal G3 domains and lacks the GAG domains (Fig. 1A)), recombinant G1 domain, or recombinant G3 domain. Note that the native versican was isolated from bovine liver and likely represents a heterogeneous mixture of versican isoforms; dot blotting confirms the presence of the GAG domains in the mixture. We found that native versican and the V3 isoform interacted with collagen in a dose-dependent manner (Figure 2A and Figure S1) and that the interaction between native versican and collagen was independent of GAGs (Figure 2B). These interactions were sensitive to pH and ionic strength (Figure S2). Given that V3, which includes only the G1 and G3 domains, bound collagen at high affinity, we tested these domains individually and found that binding activity was localized primarily to the G3 domain (Figure 2C), with minimal binding to the G1 domain. Thus, collagen and versican physically interact through the versican G3 domain.

**Figure 2.**
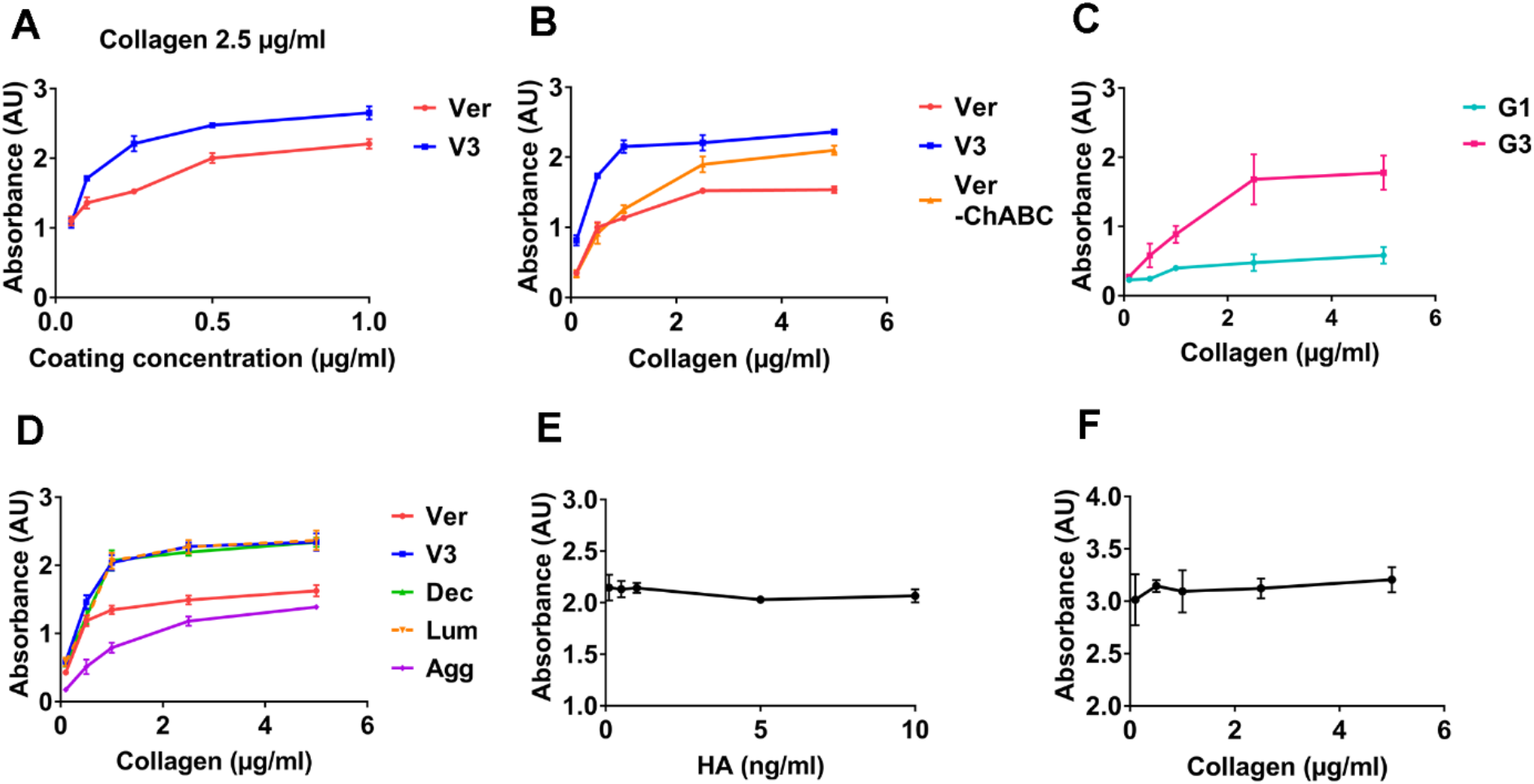
Versican interacts with collagen via its G3 domain. **A)** Plates were coated with versican (Ver, red) or the V3 isoform (V3, blue) at 0.05, 0.1, 0.25, 0.5 and 1 μg/ml. The absorbance of collagen, added at 2.5 μg/ml and detected via a biotin-conjugated antibody, was measured colorimetrically. **B)** Plates were coated with 0.25 μg/ml versican (red), V3 (blue) or versican after chondroitinase ABC digestion (Ver-ChABC, orange). The binding of increasing concentrations of collagen (0.1, 0.5, 1.0, 2.5 and 5 μg/ml) was assayed. **C)** Plates were coated with 0.25 μg/ml recombinant G1 (teal) or G3 (pink) and the binding of increasing concentrations of collagen (0.1, 0.5, 1.0, 2.5 and 5 μg/ml) was assayed. **D)** Plates were coated with 0.25 μg/ml versican (red), V3 (blue), decorin (Dec, green), lumican (Lum, orange dash line) or aggrecan (Agg, purple) and the binding of increasing concentrations of collagen (0.1, 0.5, 1.0, 2.5 and 5 μg/ml) was measured. **E)** Plates were coated with 0.25 μg/ml V3 to which was added collagen (1 μg/ml) mixed with increasing concentrations of HA (0.1, 0.5, 1, 5 and 10 ng/ml). **F)** Plates were coated with 0.25 μg/ml V3 to which was added HA (10 mg/ml) mixed with increasing concentrations of collagen (0.1, 0.5, 1.0, 2.5 and 5 μg/ml). Three independent experiments were carried out for each condition; each line in an individual graph is from the same trio of experiments. Data represent mean ± SD.

Interactions between collagen and versican, V3, and the versican core protein fitted linearly on a Scatchard plot (R^2^>0.9 for all; Table 1 and Figure S3), in contrast to interactions between collagen and G1 or G3. The maximum absorbance (ΔA_max_) also supported the identification of the G3 domain as the collagen binding site, although the dissociation constants (K_d_) showed that the binding affinity between collagen and V3 was higher than that between collagen and G3. We carried out similar assays using decorin, lumican and aggrecan (Figure 2D) and confirmed the previously-reported binding between collagen and SLRPs as well as the validity of our binding assays.

**Table 1.** The dissociation constant (K_d_) and maximum absorbance (X-intercept) analyzed by the Scatchard equation: ΔA/C=ΔA_max_/K_d_-ΔA/K_d_ (ΔA – measured value of absorbance, C – collagen concentration). By plotting ΔA/C versus ΔA, the K_d_ and ΔA_max_ were calculated from the slope and the X-intercept of the linear fitting of the Scatchard plot (Figure S3). R^2^ showed the quality of linear fitting. Three independent experiments were carried out for each; data represent mean ± SD.

| Collagen binding with | $K_d$ (nM) | $\Delta A_{max}$ | $R^2$ |
| --- | --- | --- | --- |
| Versican | $1.3 \pm 0.1$ | $1.7 \pm 0.1$ | $0.95 \pm 0.04$ |
| V3 isoform | $0.7 \pm 0.1$ | $2.5 \pm 0.1$ | $0.97 \pm 0.02$ |
| Versican digested with ChABC | $2.2 \pm 0.6$ | $2.3 \pm 0.1$ | $0.92 \pm 0.04$ |
| G1 domain | $0.9 \pm 0.4$ | $0.6 \pm 0.1$ | $0.47 \pm 0.05$ |
| G3 domain | $3.2 \pm 0.6$ | $2.2 \pm 0.3$ | $0.60 \pm 0.11$ |

Versican binds HA via its G1 domain [23] and we therefore carried out competition binding assays to confirm that versican binds collagen and HA at different locations (G3 versus G1). As shown in Figure 2E, adding HA had no impact on the collagen-V3 interaction; similarly, adding collagen did not alter the HA-V3 interaction (Figure 2F). Thus, versican can bind collagen via its C-terminal G3 domain and bind HA via its N-terminal G1 domain.

### The Collagen Toolkit identifies collagen binding sites for the V3 isoform and G3 domain

To identify versican binding sites on collagen, we used the Collagen Toolkit II [15], a library of synthetic type II collagen peptides covering the entire triple-helical region. (Table S1 shows the amino acid sequence of each Toolkit peptide; Toolkit II peptide sequences are similar to those for type I collagen.) Using a solid-phase binding assay and comparing the V3-binding capacity of each Toolkit peptide to that of full-length collagen, we showed relatively high binding of V3 to peptides II-1, II-4, II-8, II-11, II-15 and II-18 (Figure 3A). We also tested the recombinant G3 domain and found that it bound to a different set of peptides II-5 and II-44 (Figure 3B). Aligning the sequences of these positive Toolkit peptides suggests a common R-G-Hydrophobic-O motif. GPA triplets following this motif in some peptides represent a second or extended binding motif (Figure 3C). When we tested variants of peptides III-5 (which is similar to II-5) and II-44 (Figure S4), however, the binding motifs appeared to be different than (or in addition to) the R-G-Hydrophobic-O motif. The peptide II-5 derivative containing a KGHR motif, which is the conserved motif in collagen crosslink sites [16], and the peptide II-44 derivatives containing GLAGQRGIVGLOGQR both showed increased binding to G3 (Figure S4).

**Figure 3.**
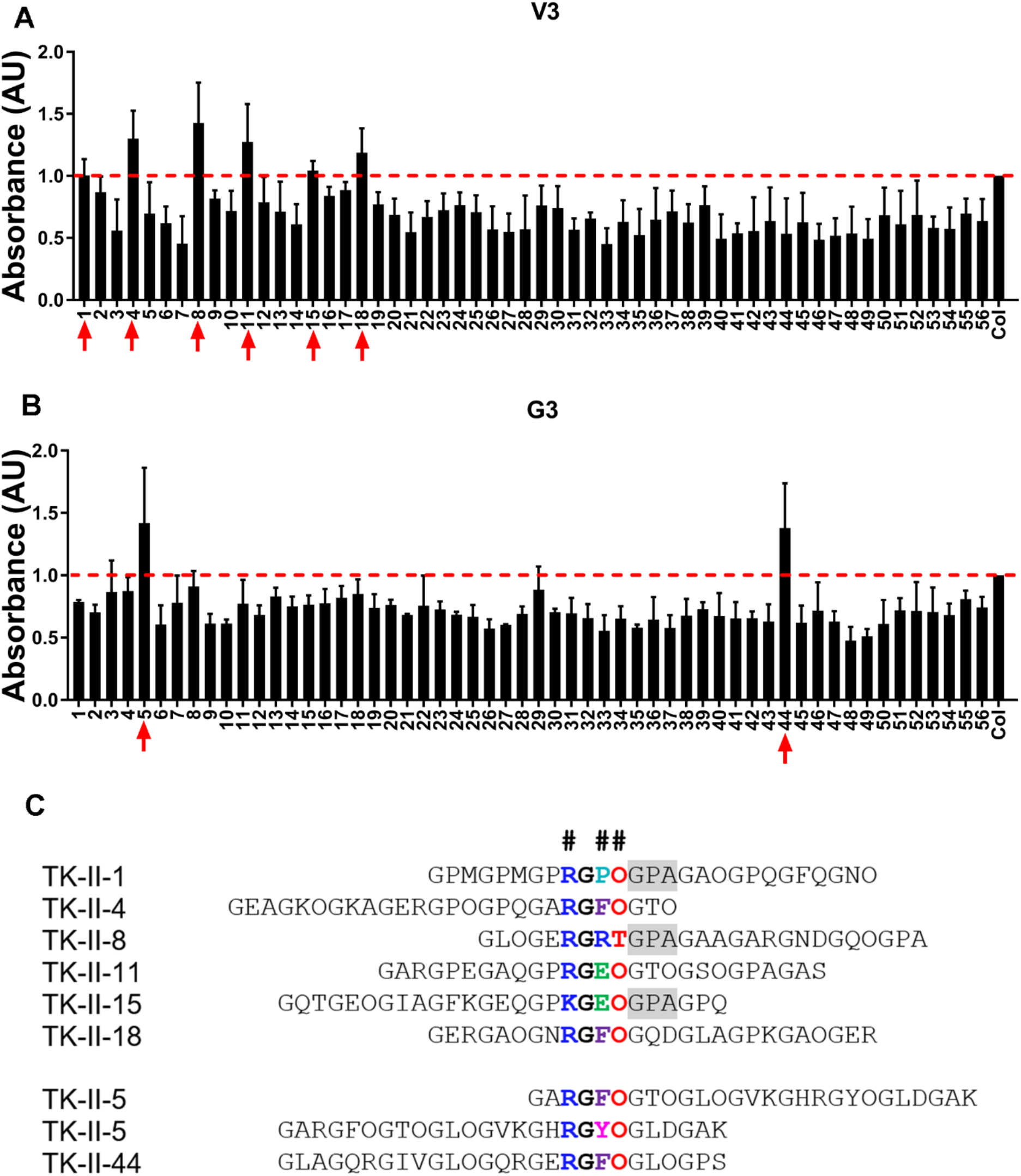
Potential versican binding sites on collagen identified using the Collagen Toolkit II. **A, B)** Binding between Collagen Toolkit II peptides and 10 μg/ml recombinant V3 (A) or G3 (B) was tested using a solid-phase binding assay. Empty wells in the Toolkit peptide plate were coated with full-length collagen as a positive control and data were normalized to the positive control. Three independent experiments were carried out. Data represent mean ± SD. **C)** Binding motifs (in color, and shaded) identified by the alignment of versican-binding Toolkit peptides. Note that peptide II-5 has two instances of the potential motif.

### Versican regulates the organization of fibroblast-derived collagen matrices

Having shown that versican binds collagen directly, we sought to define its role in matrix architecture. We used a fibroblast-derived matrix assay whereby fibroblasts were plated on dishes coated with versican isoforms, and the organization and alignment of the matrix deposited by the fibroblasts was investigated by immunostaining and collagen second harmonic generation imaging (SHG). As shown in Figure 4A-C, fibroblasts produced collagen with networked and locally highly-aligned fibers. The presence of versican or V3 led to increased SHG signal intensity (Figure 4D) and increased fiber alignment (Figure 4E). Fibronectin, a cell-associated protein that interacts with collagen, was highly expressed (surrounding each cell) for all groups (Figure 4F-H) and slightly increased for V3 (Figure 4I). Importantly, fibroblasts remained quiescent and there was no change in cell number, indicating that proliferation and myofibroblastic differentiation were not the cause of the increased collagen deposition (Figure 4J and Figure S5). Thus, versican promoted the formation of collagen-rich matrices with highly aligned fibers.

**Figure 4.**
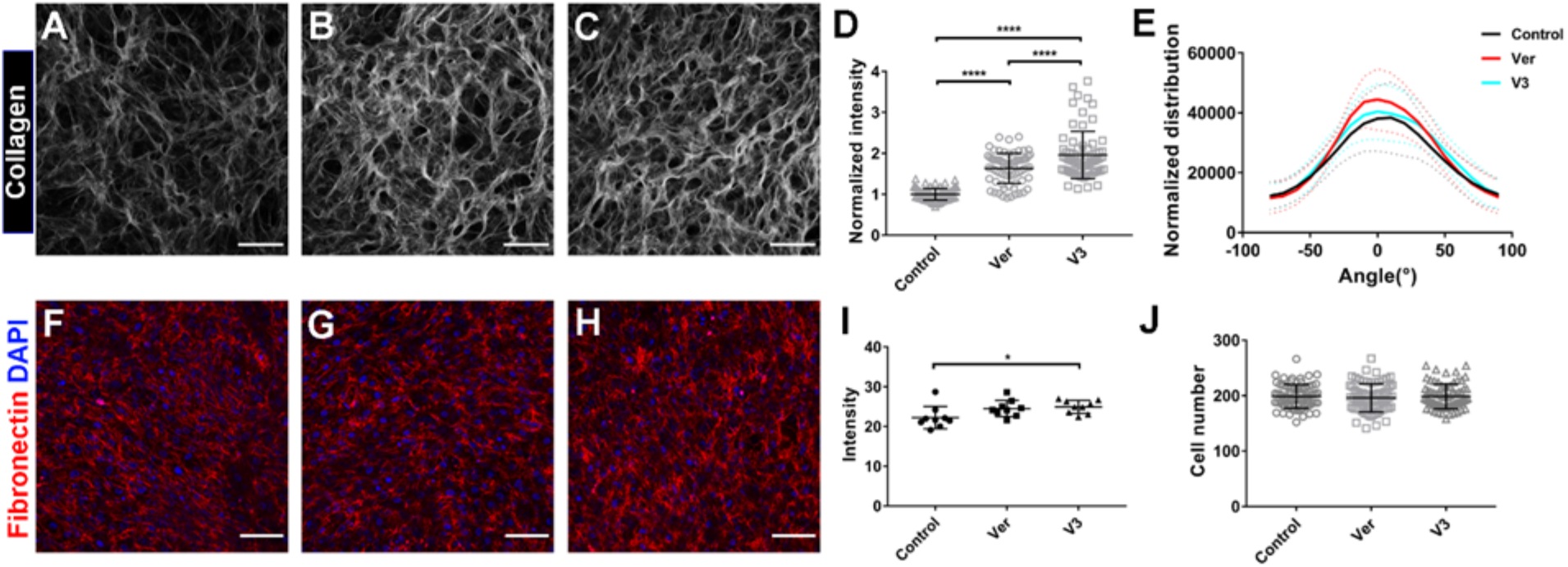
Versican and its V3 isoform upregulated the deposition of collagen-rich matrices by fibroblasts and led to increased fiber alignment. **A-C)** Representative SHG imaging of the fibroblast-deposited matrices: control plate, with vitronectin coating (A), versican-coated plate (B), V3-coated plate (C). **D)** Quantification of the intensity of the SHG signals, normalized to the value for the control group. **E)** The distribution of collagen fiber orientation was analyzed using OrientationJ. The data were normalized to the dominant angle of each SHG image. The statistical significance of differences between conditions is shown in Table S2. **F-H)** Representative confocal imaging of immunostaining for fibronectin in fibroblast-deposited matrices (red - fibronectin, blue - DAPI): vitronectin coated plate as a control (F), versican coated plate (G), V3 coated plate (H). **I)** Quantification of the intensity of fibronectin staining. **J)** Quantification of cell numbers after 7 days in culture; each data point is from 9 images of each technical repeat. Four independent experiments were carried out with two technical repeats for each coating condition per experiment. Scale bar = 100 μm. Data represent mean ± SD; D, I and J were analyzed using one-way ANOVA, E was analyzed using two-way ANOVA with repeated measurements; *P<0.5, ****P<0.0001.

### Versican has different effects on the mechanics of collagen matrices *in vitro* compared to other proteoglycans

The demonstration that versican alters the structure of the ECM [5] suggests that it could also influence its mechanics. We used shear rheometry to measure the viscoelasticity of collagen that was co-gelled with different matrix proteoglycans at the same weight ratio (Col:PG = 15:1). Gels were formed on the rheometer at 37°C, and the shear storage and shear loss moduli (G’ and G”) were measured during gelation. The addition of versican and V3 accelerated collagen gelation while neither aggrecan nor decorin caused significant changes (Figure 5A). Once gelation was complete, the presence of versican or V3 was associated with significantly decreased shear modulus (G’) compared to collagen-only gels (Figure 5B). Aggrecan, which like versican is a hyalectan and bottlebrush proteoglycan, had no impact on the G’ of collagen gels. Decorin, a SLRP, also had no effect (Figure 5B). Collagen gels that included HA, a versican binding partner, led to increased G’ compared to collagen-only gels; the presence of HA in collagen-V3 co-gels prevented the decrease in G’ observed with just collagen and V3, and the resulting collagen-V3-HA gels had similar G’ values as gels with collagen alone (Figure 5C). When we re-evaluated previously published scanning electron microscopy images of collagen and collagen-versican gels [5], we found that versican decreases total fiber length, decreases mean pore size and the number of fiber-fiber intersections (Figure S6); the decrease in collagen G’ in the presence of versican is consistent with this decreasing number of network connections [24].

**Figure 5.**
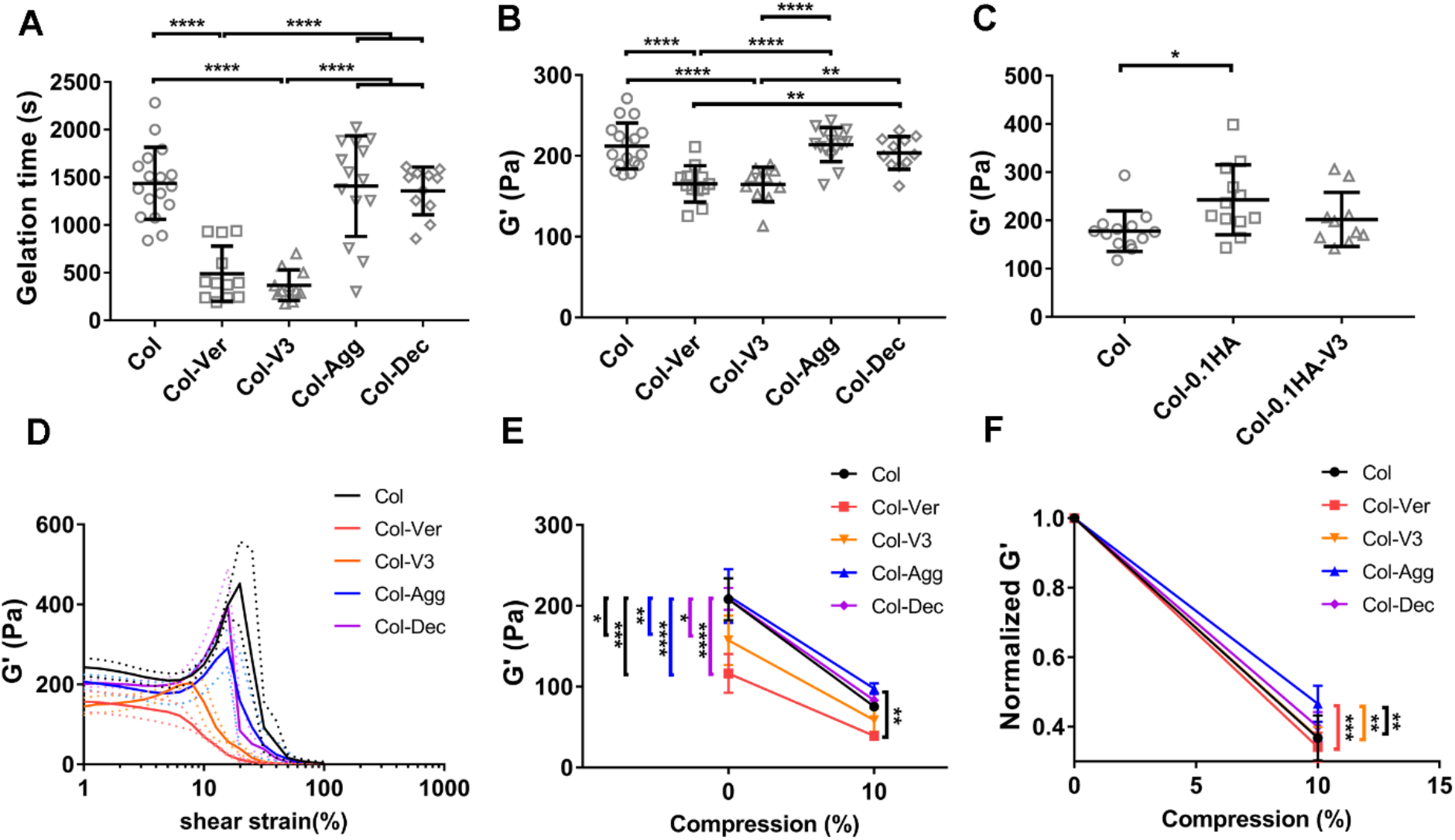
Different matrix proteoglycans have distinct effects on the mechanics of collagen networks. **A)** Gelation times for collagen-proteoglycan co-gels, with rheological measurements taken during gelation. Col: 2.5 mg/ml pure collagen gel; Col-Ver: 2.5 mg/ml collagen with 0.167 mg/ml versican; Col-V3: 2.5 mg/ml collagen gel with 0.167 mg/ml V3; Col-Agg: 2.5 mg/ml collagen gel with 0.167 mg/ml aggrecan; Col-Dec: 2.5 mg/ml collagen gel with 0.167 mg/ml decorin. **B)** The shear storage modulus (G’) for the gels in (A), measured after complete gelation. **C)** G’ for collagen gelled with addition of high molecular weight (1.5 MDa) HA or both HA and V3. Col-0.1HA: 2.5 mg/ml collagen gel containing 0.1 mg/ml HA; Col-0.1HA-V3: 2.5 mg/ml collagen gel containing 0.1 mg/ml HA and 0.167 mg/ml V3. **D)** G’ of different collagen-proteoglycan co-gels measured by shear rheometry under increasing strain stiffening after full gelation (see Table S3). **E)** G’ of different collagen-proteoglycan co-gels measured with the gap remaining at 1 mm and under compression to 10% (gap changed to 0.9 mm). **F)** G’ values at 10% compression for the different co-gels in (E) were normalized to the G’ in the non-compressed state. For measuring G’ during gelation, N=17 for Col, N=12 for Col-Ver, N=11 for Col-V3, N=15 for Col-Agg and N=11 for Col-Dec. These gels were tested with either strain sweep (N=3 for Col, N=3 for Col-Ver, N=3 for Col-V3, N=4 for Col-Agg and N=3 for Col-Dec) or compression (N=3 for Col, N=3 for Col-Ver, N=4 for Col-V3, N=4 for Col-Agg and N=3 for Col-Dec); each gel was subject to only one test. The dotted lines in (D) represent SD. Data represent mean ± SD; A, B, C, and G were analyzed using one-way ANOVA, D, E, and F were analyzed using two-way ANOVA with repeated measurements; *P<0.5, **P<0.01, ***P<0.001, ****P<0.0001.

Isolated collagen has complex non-linear elastic behaviors including significant shear strain stiffening and compression softening [25]. The addition of versican eliminated the strain stiffening behavior of collagen and the addition of V3 led to markedly blunted strain stiffening (Figure 5D, compare red and orange curves to control curve in black; Table S3). Collagen cogelled with aggrecan or decorin underwent strain stiffening, but slightly less than pure collagen (Figure 5D). The strain at which collagen-only gels failed was higher than for collagen gels with added V3, aggrecan, or decorin. Collagen gels, either without additions or co-gelled with versican, V3, aggrecan, or decorin, showed compression-softening behaviors (Figure 5E, F). The addition of aggrecan significantly attenuated compression softening in contrast to other conditions (Figure 5F blue curve). Thus, overall, versican and the V3 isoform have a markedly different impact on collagen mechanics than do other proteoglycans.

### Versican and its GAG side chains play a role in tissue mechanics

To test the impact of versican on collagen organization and mechanics *in vivo*, we used liver, the tissue mechanics of which have been well-characterized using shear rheometry [26]. We first perfused rat liver in situ with ADAMTS-5 or chondroitinase ABC (ChABC); the former cleaves versican in the β-GAG domain [27][28] and the latter removes the chondroitin sulfate chains. Although ADAMTS5 cleaves hyalectan family members generally [29] and neither enzyme is specific for versican, versican is the major hyalectan and CS-attached protein in liver, and is the major target of both enzymes in this tissue [30][31]. To test the effectiveness of our enzymatic perfusions, sulfated GAGs in perfused liver tissues were quantified and showed a significant decrease after both ADAMTS-5 and ChABC perfusion (Figure 6A). There are numerous ADAMTS-5 cleavage sites on versican, potentially leading to the production of numerous small fragments; the data suggest they diffused out of the liver during perfusion [32]. Immunostaining using an anti-DPEAAE antibody, which recognizes the epitope exposed by ADAMTS-5 cleavage of versican, was positive only for ADAMTS-5 perfused liver, and staining by an antibody that recognizes only non-cleaved versican was significantly decreased (Figure 6B and Figure S7).

**Figure 6.**
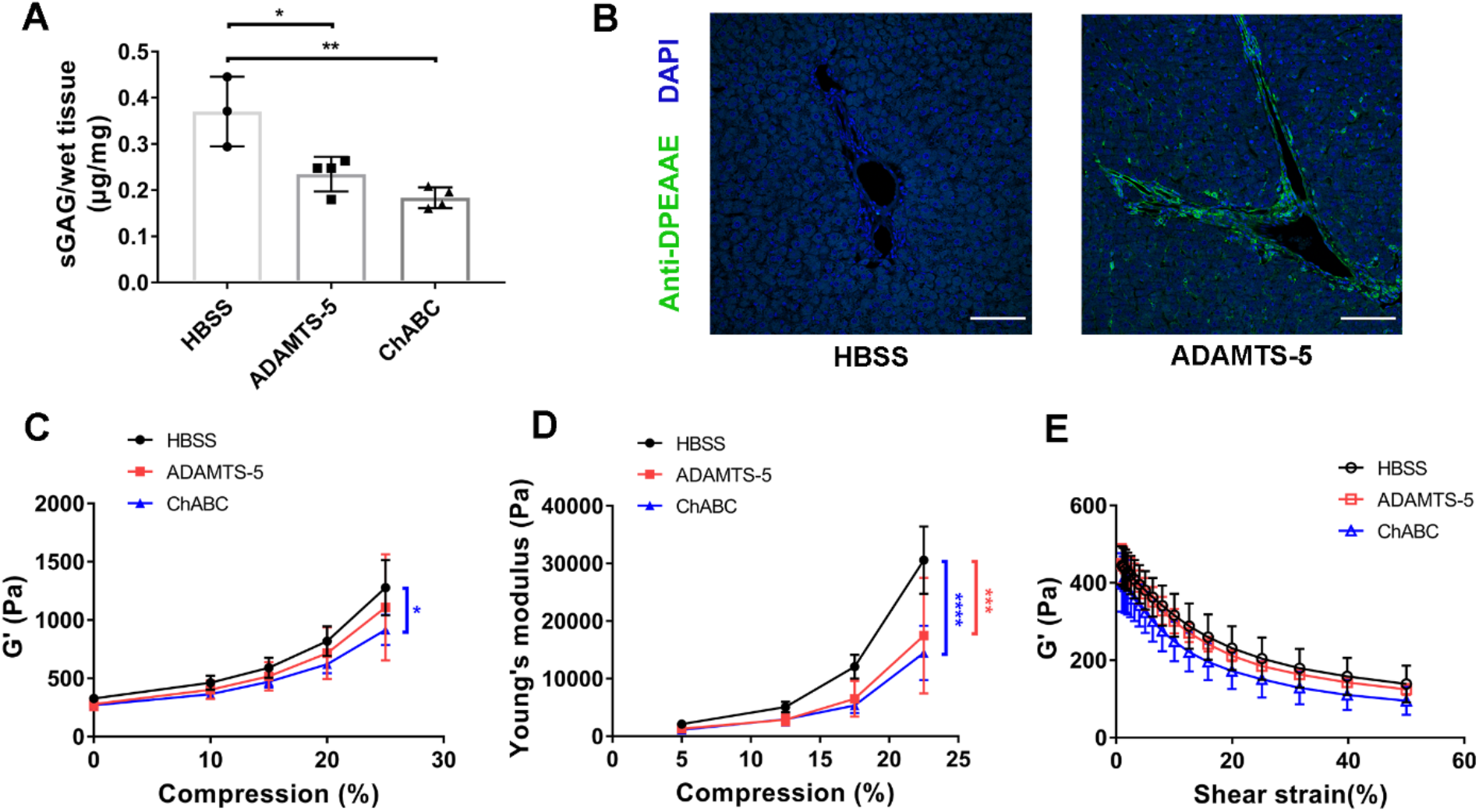
ADAMTS-5 and chondroitinase ABC treatment of rat livers alters compression stiffening behavior. **A)** Sulfated GAG quantification after perfusion with either enzyme. **B)** Representative confocal imaging of immunostaining using an antibody against DPEAAE, the epitope exposed by ADAMTS-5 cleavage of versican. DPEAAE (green), DAPI (blue). **C)** G’ was measured under 0%, 10%, 15%, 20% and 25% compression (with HBSS perfusion as a control). There is a significant difference at 25% compression. **D)** Young’s modulus (*E*) was calculated from normal force and gap changes and plotted at 5%, 12.5%, 17.5% and 22.5% compression (significance shown for 22.5% compression). **E)** G’ measured at increasing strain from 1% to 50%. N=3 for HBSS, N=4 for ADAMTS-5 and for ChABC perfused livers. Data did not show statistical differences. Compression and strain sweep experiments were done on the same liver samples, as was the assay for sulfated GAGs (A). Scale bar = 200 μm. Data represent mean ± SD; A was analyzed using one-way ANOVA, C, D, and E using two-way ANOVA with repeated measurements; *P<0.5, **P<0.01, ***P<0.001, ****P<0.0001.

We first examined G’ and compression stiffening. While biopolymers like collagen compression soften, tissues undergo compression stiffening [33], and this was attenuated by perfusion with either enzyme. The removal of chondroitin sulfate caused a significant decrease in G’ under 25% compression (Figure 6C), as was previously reported after the more global removal of GAGs using α-amylase [26]. There were dramatic changes in compression stiffening measured by Young’s modulus (*E*), with both enzymatic perfusions resulting in significantly decreased *E* at 22.5% compression (Figure 6D). Liver normally undergoes shear strain softening; this was not significantly affected by perfusion with either ADAMTS-5 or ChABC (Figure 6E). Thus, perfusion with either ChABC or ADAMTS-5, which primarily affects versican in liver tissue, significantly alters liver mechanics.

## Discussion

Matrix proteoglycans are important collagen-binding ECM proteins and key regulators of collagen behaviors. We report here that versican, a widely-distributed large chondroitin sulfate proteoglycan, binds to and regulates the mechanics of type I collagen. The binding of versican to collagen is through the C-terminal G3 domain of its core protein, independent of its GAG modifications, and distant from the versican-HA binding site (Figure 7A), and the impact of versican on collagen mechanics is significant and different than for other matrix proteoglycans.

**Figure 7.**
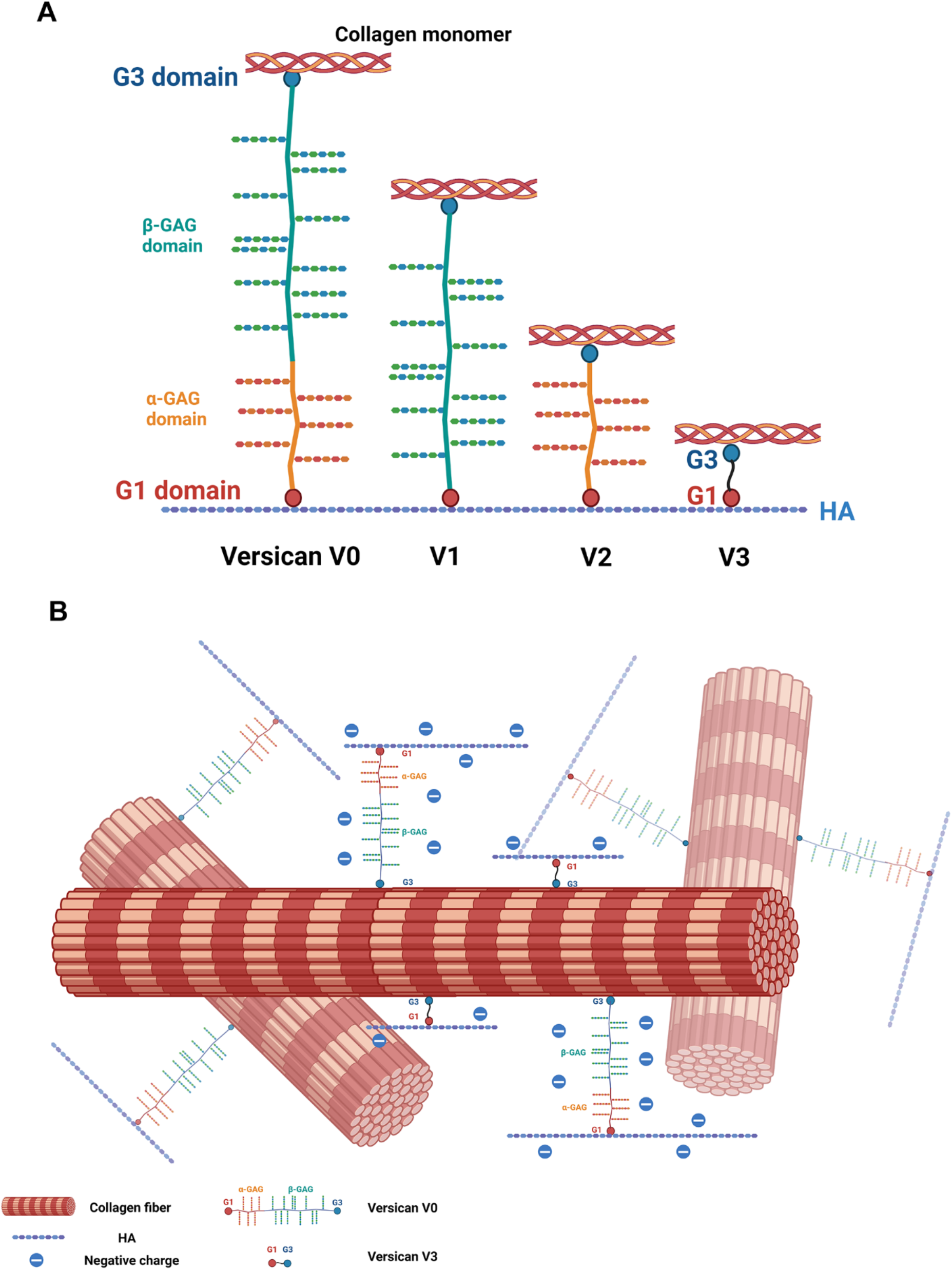
Model of interactions between HA, collagen, and versican. **A)** Collagen and HA binding sites on different versican isoforms. **B)** Versican may serve as a linker between collagen fibers and HA chains in collagen fibrous networks.

Solid-phase binding assays show binding between collagen and versican, V3, and G3, but not G1. Although the full-length versican used was isolated from tissue and contains a variety of isoforms as well as small amounts of other proteins, the binding to recombinant V3 and the similarity between the binding curves suggests that the recorded binding is to versican itself. The binding between collagen and versican at the versican G3 domain was confirmed by use of the Collagen Toolkit II, a reagent set based on the sequence of type II collagen but well-established as a reporter of binding to type I collagen (Figure 3, Table S1) [15]. The peptides to which V3 binds (II-1, II-4, II-8, II-11, II-15, and II-18) are, with the exception of II-8, unusual binding sites and hint at as yet unknown functions for the interaction between collagen and V3 (and potentially full-length versican). The use of the G3 domain to probe the Toolkit II showed that its binding motifs were different than for V3 (peptides II-5 and II-44). This raises the possibility that the two domains (G1 and G3) that make up V3 are folded in such a way that key collagen binding sites are masked or uncovered by the interaction between the domains. Whether this would similarly be true for full-length versican (V0 or V1) is not known. Unfortunately, long-form isoforms of versican are not available at sufficient purity to test with the Toolkit, although there is the possibility that they would demonstrate additional binding sites not observed with V3 alone. The binding sites identified by G3 are peptide II-5, incorporating a crosslinking site (KGHR), and peptide II-44, a collagenase cleavage site [15][16]. Using a similar Toolkit assay, fibromodulin was found to interact with the same KGHR motif and facilitate lysyl oxidase mediated collagen crosslinking [16]. This suggests that G3 may, either as an isolated domain (if it exists in that form) or as part of the full-length protein, serve as a modulator of collagen fiber assembly and degradation.

Alignment of the positive binding peptides of V3 and G3 identifies an R-G-Hydrophobic-O motif, in some cases with an adjacent GPA triplet. In the base motif, R is substituted by K in Toolkit II-15 and the aliphatic stems of R, E and P and the aromatic rings of F and Y provide hydrophobic interactions. The R-G-Hydrophobic-O motifs are found in other Toolkit II peptides that have not been identified in our assay as binding peptides, suggesting the potential importance of other factors including distant sequences and localization on different faces of the triple helix. For the GPA triplets, the methyl side chain of A could minimize steric hindrance but not contribute to binding directly. Defining the 3D structure of V3 and G3 will be important to better understand the structure of the collagen/versican complex.

HA was shown previously to bind versican via the G1 domain [23], and our data show that the HA and collagen binding sites are separate, although analysis of G1 binding to collagen with a Collagen Toolkit could be done in the future. This suggests that versican serves as a linker between HA chains and collagen in the ECM and that different versican isoforms could have distinct size-dependent effects in regulating the nature of the versican/HA/collagen complex (Figure 7; the diameter of the collagen fibril is roughly 200 nm and the length of isolated versican is similar). More generally, versican and its isoforms could regulate the fibrous network of the ECM, particularly the relationship between its structural and space-filling properties (Figure 7). The structurally-similar bottlebrush proteoglycan aggrecan binds to collagen via its keratan sulfate domain, close to the G1 domain HA binding site, suggesting that the interactions between collagen and versican and between collagen and aggrecan are different and that the two proteoglycans have different linker functions. This is consistent with the differences we observe in collagen gel mechanics when either versican or aggrecan is added. Notably, the K_d_ is 1.1 μM for binding between collagen and aggrecan keratan sulfate domain [18], while the K_d_ for binding between collagen and versican is approximately 1.3 nM.

We previously reported that versican and V3 have different effects on collagen fibrillogenesis and organization than aggrecan and the two SLRPs decorin and lumican. This is interesting given that mice deficient in many of the SLRPs have abnormal collagen fibril development [10][34][35], and raises questions about the role of versican in regulating collagen deposition and organization. As an example, immunostaining of mouse extrahepatic bile ducts shows a submucosa at day of life 0 rich in HA and versican, with few if any collagen fibrils; collagen is deposited into this GAG-filled milieu in the first two weeks of life [22]. This, combined with IEM images (Figure 1B-D) demonstrating co-localization between collagen and versican at day of life 3, raises the interesting possibility that versican/V3 (in combination with lumican and decorin, which are also present in the neonatal duct [22]) potentially regulate collagen deposition and organization in this tissue. Importantly, the K_d_s for binding between collagen and versican, collagen and fibromodulin, and collagen and decorin are similar at 1.3 nM (Table 1), 9.9 nM [36] and 14 nM [37], respectively. The pattern of interaction between versican and collagen seen on IEM is quite different, however, from the pattern observed with decorin, which is arranged in ring or spiral-like structures along collagen fibrils [38].

Fibroblast-derived matrix studies demonstrate that the presence of versican or V3 increases both the production and the alignment of collagen by fibroblasts (Figure 4). Given that collagen architecture plays a key role in fibroproliferative processes such as fibrosis and metastasis [39][40], this finding suggests that versican might play a role in pathology as well as normal wound healing. For example, versican expression and its cleavage via ADAMTS-1, 4, 5, 8, 9, 15, 20 are dysregulated during liver fibrosis and recovery [20]. The versican-mediated alterations in tissue G’ and compression stiffening we report are potentially similarly important to a variety of pathologies.

We studied the impact of versican on collagen mechanics using shear rheometry. By measuring the G’ of collagen gels and co-gels during gelation, we observed that the addition of versican or V3 accelerated gelation (consistent with our data using an *in vitro* turbidity assay [5]) and led to a decrease in G’ of the final gel. As noted above, the decrease in G’ is consistent with a decreasing number of network connections (Figure S6) [24]. Although versican has significant GAG modifications that might be expected to impact mechanics in collagen-versican co-gels, it may be that the overall difference in network organization has a more significant effect on mechanics than GAGs.

To investigate the effect of versican on native tissue mechanics, we perfused liver with ADAMTS-5 and with ChABC, and studied its mechanics using shear rheometry. Notably, although neither enzyme is specific for versican, versican is the major target in liver for both. ADAMTS-5 cuts versican at a well-studied cleavage site at Glu441-Ala442 (for V1) [27], but there is also evidence for multiple ADAMTS-5 cleavage sites on the versican core protein [32][41], making it likely that the enzyme digests versican into small core protein fragments. Consistent with this, ADAMTS-5 perfusion led to significant loss of GAGs from the tissue, suggesting that small GAG-attached core protein fragments were generated by ADAMTS-5 and flushed out during continued tissue perfusion. Although we found that the inclusion of versican into collagen gels decreased gel stiffness (Figure 5B), disruption of versican and its GAGs had no impact on baseline liver stiffness but eliminated compression stiffening in tissue (Figure 6). The difference between *in vitro* and *in vivo* results is likely related to the fact that versican and collagen in tissue are in a preformed network; the enzymatic digestions we performed were unlikely to alter collagen cross-linking and network structure, and the effects on GAGs predominated. This is consistent with our previously-reported finding that perfusion with α-amylase, which digests α-linked polysaccharides such as GAGs, eliminated compression stiffening without significantly impacting baseline G’.

## Conclusion

In sum, we demonstrate a direct binding interaction between collagen and versican and report that versican modulates collagen organization and mechanics. Versican binds collagen and HA through different and, in the full-length protein, widely-separated domains, suggesting that it serves as a linker between the two and may be important in regulating both structural and space-filling functions of the ECM. Versican has a significant impact on architecture and mechanics of collagenous matrices, and is thus a potential regulator of a number of pathological processes including fibrosis and metastasis. Further investigation of the structure, isoform kinetics, and deposition pattern of versican would contribute to understanding the mechanism of the structural and mechanical alterations that occur during fibroproliferative and other diseases.

## Methods

### Animal Studies

All animal studies followed the Guide for the Care and Use of Laboratory Animals of the National Institutes of Health. The animal protocol (#804031) was approved by the Institutional Animal Care and Use Committee of the University of Pennsylvania. 300-350 g Sprague-Dawley rats (Charles River Laboratories, Malvern, PA) were housed in pairs, fed standard chow, and exposed to 12 hour light-dark cycles.

### Reagents and cells

Rat tail type I telo-collagen, used for binding assays and immune-electron microscopy, was from Corning (Corning, NY, USA). Type I atelo-collagen from calf skin, used for rheometry, was purchased from MP Biomedicals (Irvine, CA, USA). Versican was isolated from bovine liver [5]. Aggrecan and decorin, both isolated from bovine cartilage, were purchased from Sigma (St. Louis, MO, USA). Recombinant human versican isoform V3 and lumican were from R&D Systems (Minneapolis, MN, USA). Recombinant human versican protein G1 and G3 domains (ab152303 and ab236178) were from Abcam. Biotinylated versican G1 domain was from Echelon Biosciences (Salt Lake City, UT, USA). Sodium hyaluronan (1.5MDa) was from Lifecore (Chaska, MN, USA) and hyaluronan biotin sodium salt was from Sigma. Casein blocking buffer (10×) was from Sigma. The Collagen Toolkit was from CambCol Laboratories (Ely, UK). High-sensitivity streptavidin-horseradish peroxidase (HRP) and tetramethylbenzidine (TMB) were from Thermo Fisher Scientific (Waltham, MA, USA). Gelatin from porcine skin and ethanolamine were from Sigma. Glutaraldehyde solution (50%) was purchased from Fisher Chemicals. Dulbecco’s phosphate buffered saline with calcium chloride and magnesium chloride (DPBS+ (10×)) and Dulbecco’s phosphate buffered saline without calcium chloride and magnesium chloride (DBPS- (1×)) were from Life Technologies. Vitronectin (recombinant human protein) was from Fisher Scientific.

NIH 3T3 fibroblasts were obtained from the ATCC (Manassas, VA, USA). They were cultured in Dulbecco’s Modification of Eagle’s Medium (DMEM) with 4.5 g/L glucose and L-glutamine without sodium pyruvate (Corning) supplemented with 10% calf serum (Thermo Fisher), 1% penicillin/streptomycin (Corning) and 0.5% fungizone (Life Technologies, Carlsbad, CA, USA) at 37°C in a humidified atmosphere with 5% CO_2_/balance air.

### Immunoelectron microscopy

Extrahepatic bile ducts were dissected from neonatal (day 3) and adult mice, then were fixed with 4.0% paraformaldehyde and 0.1% glutaraldehyde in 0.1M sodium cacodylate buffer (pH 7.4) at 4°C overnight. Fixed samples were washed with buffer and rinsed in diH_2_O, then dehydrated through a graded ethanol series and embedded in LRWhite (London Resin Company, Berkshire, England). Thin sections were stained with primary anti-versican (ab1033; Sigma) antibody at 1:10 at 4°C overnight. After rinsing, sections were incubated with secondary anti-rabbit antibody (ab105295; Abcam; 12nm nanogold conjugated) at 1:50 at 4°C overnight. After rinsing, sections were stained with phosphotungstatic acid (Electron Microscopy Science, Fort Washington, USA) and uranyl acetate (Electron Microscopy Science, Fort Washington, USA). Sections were examined with a JEOL 1010 electron microscope fitted with a Hamamatsu digital camera and AMT Advantage NanoSprint500 software. Collagen fibers with gold particles attached were captured at 75,000×. To quantify versican-collagen colocalization, the gold particles overlying collagen fibers were counted and divided by the total fiber area, defined using ImageJ.

For studying the localization of versican in collagen gels *in vitro*, both pure collagen (as a control) and collagen-versican mixtures (weight ratio = 15:1) were prepared by diluting and neutralizing collagen to a final concentration of 1.5 mg/ml. These were gelled at the bottom of a 1.5 ml tube. The gels were then fixed with 10% formalin and permeated with 0.1% Triton-X. After rinsing, the gels were stained as described for bile duct samples.

### Solid phase binding assay

Binding between collagen and versican was studied using a solid phase binding assay. A 96-well plate was incubated with isolated versican or recombinant V3 isoform at 0.05, 0.1, 0.25, 0.5 and 1 μg/ml at 4°C overnight. The plate was then blocked using 3% BSA in TTBS (tris-buffered saline with 0.05%Tween-20) or casein blocking buffer for 3h at RT. Rat tail type I collagen was added at 0.1, 0.5, 1.0, 2.5 and 5 μg/ml and incubated overnight at RT. After rinsing, collagen was detected by incubating with biotin-conjugated anti-collagen I antibody (diluted 1:1000; ab24821; Abcam) at 37°C for 1h. The plate was rinsed again and then incubated with streptavidin-HRP (diluted 1:4000) for 30 min at RT and TMB was added until color developed (10 min). 2N sulfuric acid was added to end the reaction and the absorbance was read at 450 nm. To study the effect of pH on the interaction between collagen and versican, the pH of the binding buffer was adjusted to 6.0, 7.4 and 8.0. To study the effect of ionic strength, the ionic strength of the binding buffer was adjusted by adding sodium chloride to reach a final concentration of 0.05, 0.1, 0.15, 0.2, 0.25, 0.3, 0.35, 0.4, 0.45, 0.5, 0.55, and 0.6 M [13]. To study the interaction between collagen and the versican core protein, isolated versican was incubated with 250 mU chondroitinase ABC (Sigma) per mg substrate (in 50 mM sodium acetate, pH=8.0) at 37°C overnight to remove GAG side chains. Samples were then dialyzed with distilled water to remove intact side chains. To compare the G1 and G3 domains, 0.25 μg/ml recombinant G1 or G3 was used for coating, and collagen was added at 0.1, 0.5, 1.0, 2.5 and 5 μg/ml for binding. To compare different matrix proteoglycans, versican, V3, decorin, lumican and aggrecan were coated at 0.25 μg/ml. To investigate the effect of HA on the collagen-V3 interaction, the plate was coated with V3 and bound with 1 μg/ml collagen which had been premixed with HA (200 kDa; Lifecore) to final concentrations of 0.1, 0.5, 1, 5, and 10 ng/ml. Similarly, V3-coated plates were bound with 10 ng/ml HA (biotinylated; Sigma) mixed with collagen to a final concentration of 0.1, 0.5, 1.0, 2.5 and 5 μg/ml.

Toolkit peptide-coated plates were blocked with casein blocking buffer. 10 μg/ml recombinant V3 or G3 was added and incubated overnight. After rinsing, plates were incubated with anti-His antibody (HRP) at 1:1000 and then with TMB. Absorbance was read at 450 nm.

### Fibroblast derived matrices

Fibroblast-derived matrices were generated according to a published protocol [42] with modifications as described here. MatTek glass-bottomed dishes were incubated with 0.2% (w/v) gelatin solution (in DPBS+) for 1h at 37°C. Dishes were rinsed with DPBS+, incubated with 1% (v/v) glutaraldehyde (in DPBS+) at room temperature for 30 min and then incubated with 1M ethanolamine (in diH_2_O) at room temperature for 30 min after rinsing. After further rinsing, 0.1 mg/ml versican, V3 or vitronectin (as a control) were used to coat glass-bottomed culture dishes by incubating overnight at 37°C. Semi-confluent 3T3 fibroblasts were trypsinized and seeded on the coated dishes at 2.5×10^5^ cells/ml. After culturing overnight, culture media were replaced with media containing 100 μg/ml ascorbic acid and changed every 48h. At the third media change, additional versican, V3 or vitronectin solution was added. After 7 days of culture, matrices were rinsed with DPBS-, fixed with 10% formalin and stored at 4°C. A Leica SP8-MP spectral imaging confocal/dual-photon microscope (Leica Microsystems, Inc., Mannheim, Germany) with a linear polarizer and numerical aperture of 1.0 was used to collect SHG signals from collagen fibers. The orientation of collagen fibers was analyzed using ImageJ and its plugin OrientationJ. Briefly, the dominant angle of collagen fibrils was calculated using the Orientation Dominant Direction option and was used for angle normalization. The orientation distribution of fibrils was quantified using the OrientationJ Distribution option: the σ of pixels in the Gaussian window was set to 3; Gaussian Gradient was chosen; the Minimum Coherency and Energy setting was 0%; and the following options were selected – Orientation in the Hue section, Coherency in the Saturation section and Original-image in the Brightness section. After running the analysis, the list of orientations (degrees) and the distribution of orientation was normalized to its dominant angle. The orientation distribution was plotted using GraphPad.

For immunostaining, Fibroblast-derived matrices were stained using anti-fibronectin (1:100; ab2413; Abcam) and anti α-smooth muscle actin (α-SMA, 1:100; A2547; Sigma) antibodies and then stained with Cy3 anti-rabbit secondary antibody (1:600; 111-165-003; Jackson ImmunoResearch). Fibroblast-derived matrix samples were mounted, imaged with confocal microscopy and quantified using ImageJ.

### Collagen gel rheology

Type I collagen from calf skin was diluted to a final concentration of 2.5 mg/ml in 1x PBS at a pH=7.4. Versican, V3, aggrecan or decorin were added to the collagen solution to a final concentration of 0.167 mg/ml (Col: proteoglycan weight ratio = 15:1). A shear rheometer (Kinexus) with rSpace software was used to generate the rheological data. The temperature was set to 37°C for gelation and a 20 mm plate was used for testing. 314 μL collagen solution was added between plates (gap=1 mm). Both shear storage and shear loss moduli (G’ and G”) were measured during gelation by applying an oscillatory shear strain of 2% at a frequency of 10 rad/sec. When the shear modulus reached equilibrium, indicating complete gelation, the freshly-formed collagen gel was strain sweep tested at a frequency of 1 rad/s, during which the shear strain was increased from 1% to 100% (by logarithmic progression). Other freshly-formed gels were compressed to 10% and used to measure the shear modulus under compression. Separate gels were used for strain sweep and compression experiments.

### Liver perfusion

Rats were anesthetized with pentobarbital by intraperitoneal injection (1ml per 500g). The abdomen was opened, 5 ml 1000 USP unit/ml heparin was injected and the portal vein was catheterized and flushed with warm HBSS (without Ca^2+^, at 37°C). The inferior vena cava was then transected. To enzymatically digest versican into versikine, livers were perfused with 5 μg/200 ml ADAMTS-5 (Sigma) for 1h. To digest chondroitin sulfate, livers were perfused with 5U chondroitinase ABC (Sigma) for 1h. For control groups, livers were perfused with HBSS for 1h. After perfusion, livers were harvested and the largest lobules used for rheometry. Some samples were flash frozen for GAG quantification. Some were fixed with 4% paraformaldehyde and paraffin embedded and sectioned. Other samples were frozen in OCT and sectioned. Sections were stained with anti-versican antibody ab1033 (1:100; Sigma) for versican and ab19345 (1:100; Abcam) for versikine.

### Liver rheology

Samples for rheometry were cut using a 20 mm punch. Samples were kept hydrated throughout testing. The plate-tissue contact point was set as the normal force reached 10g (equal to 0.1N). The rheological test sequence was: (1) dynamic time sweep; (2) dynamic strain sweep. During the time sweep test, the shear storage and shear loss moduli (G’ & G”) and normal force without compression were measured at 2% strain with an oscillation frequency of 1 rad/s over 120s. These measurements were then taken under increasing uniaxial compression at 10, 15, 20 and 25%, then returned to 0%, by setting the gap size. A strain sweep was done by increasing the strain amplitude from 1% to 50% (by logarithmic progression) with an oscillation frequency of 10 rad/s.

### Statistical analysis

All results were analyzed via GraphPad Prism 7 (San Diego, CA, USA) using unpaired t test, one-way or two-way ANOVA. P values were determined by Tukey’s multiple comparison test, in which *P<0.05 was considered to be statistically significant.

## Declaration of Completing Interests

Arkadiusz Bonna and Richard W. Farndale are employees of CambCol Laboratories.

## Acknowledgements

We are grateful for the assistance of the Electron Microscopy Resource Laboratory in the Department of Biochemistry & Biophysics at the University of Pennsylvania, the UPenn NIDDK Center for Molecular Studies in Digestive and Liver Diseases Molecular Pathology and Imaging Core (P30DK050306), and Gordon Ruthel and the Penn Vet Imaging Core. We appreciate the advice provided by Edna Cukierman. Figures 1A and 7 were constructed using Biorender.

## Author Contributions

DC carried out all experiments, analyzed data and wrote the manuscript. YD, JL and BZ carried out some of the experiments with DC. AB and RWF provided experimental reagents and technical information and assisted in interpretation. PAJ provided technical information, experimental design and interpretation. RGW assisted in experimental design, interpretation and funding, and wrote the manuscript. All authors reviewed and approved the final version.

