## Supplemental Material for "Versican binds collagen via its G3 domain and regulates the organization and mechanics of collagenous matrices"

### Supplementary material

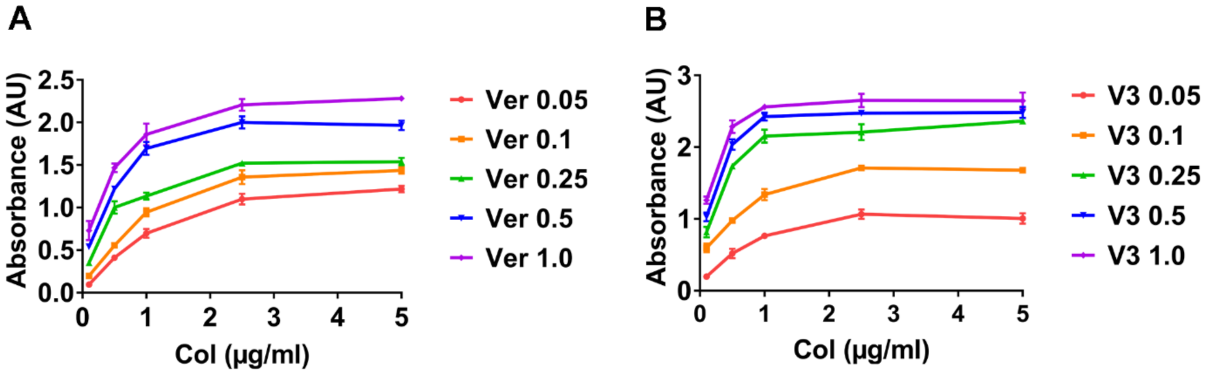


**Figure S1.** Versican-collagen binding is dose-dependent. **A, B)**The interaction between collagen and versican (A) or V3 (B) at increasing collagen and versican concentrations. A 96-well plate was coated with versican or V3 at 0.05, 0.1, 0.25, 0.5 and 1.0 µg/ml and collagen was added at 0.1, 0.5, 1.0, 2.5 and 5.0 µg/ml. The absorbance values represent the amount of collagen bound to versican or V3. Three independent experiments were carried out for each; data represent mean ± SD.

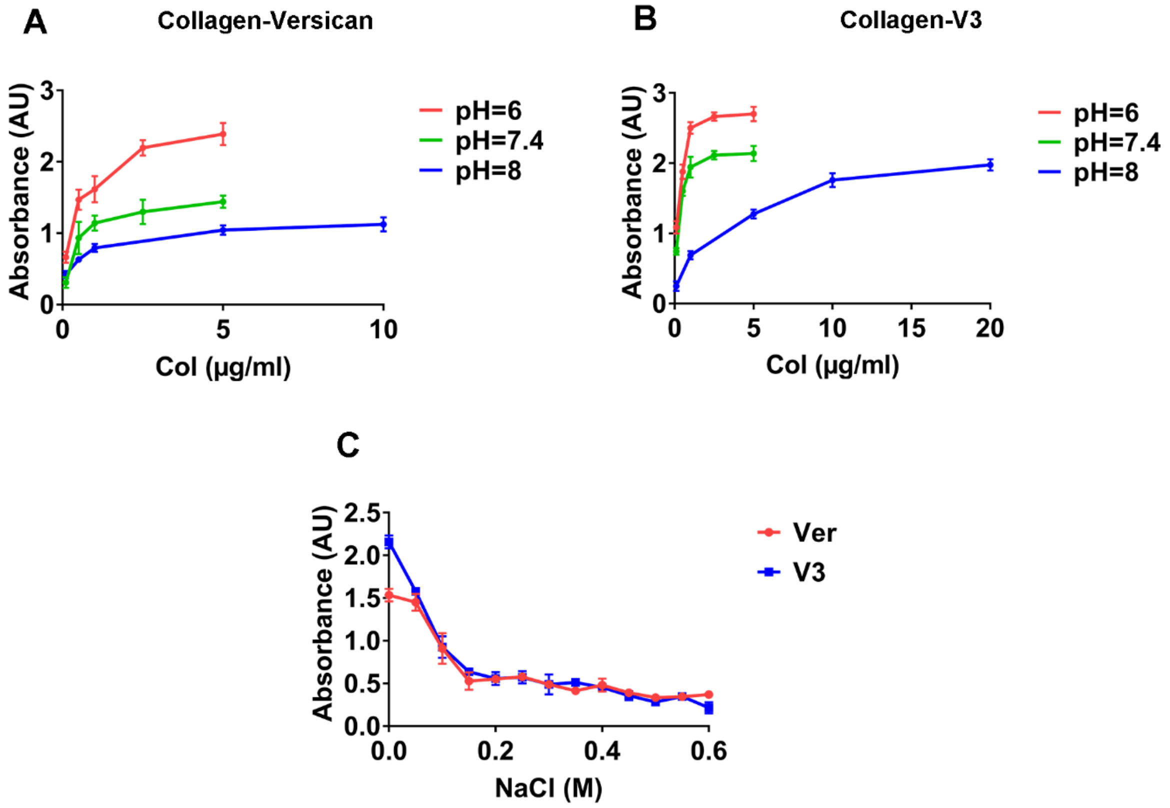


**Figure S2.** The interaction between collagen and versican is pH and ionic strength dependent. **A)** The effect of pH on collagen-versican interactions. **B)** The effect of pH on collagen-V3 interactions. **C)** The effect of ionic strength on collagen-versican and collagen-V3 interactions. Three independent experiments were carried out for each; data represent mean ± SD.

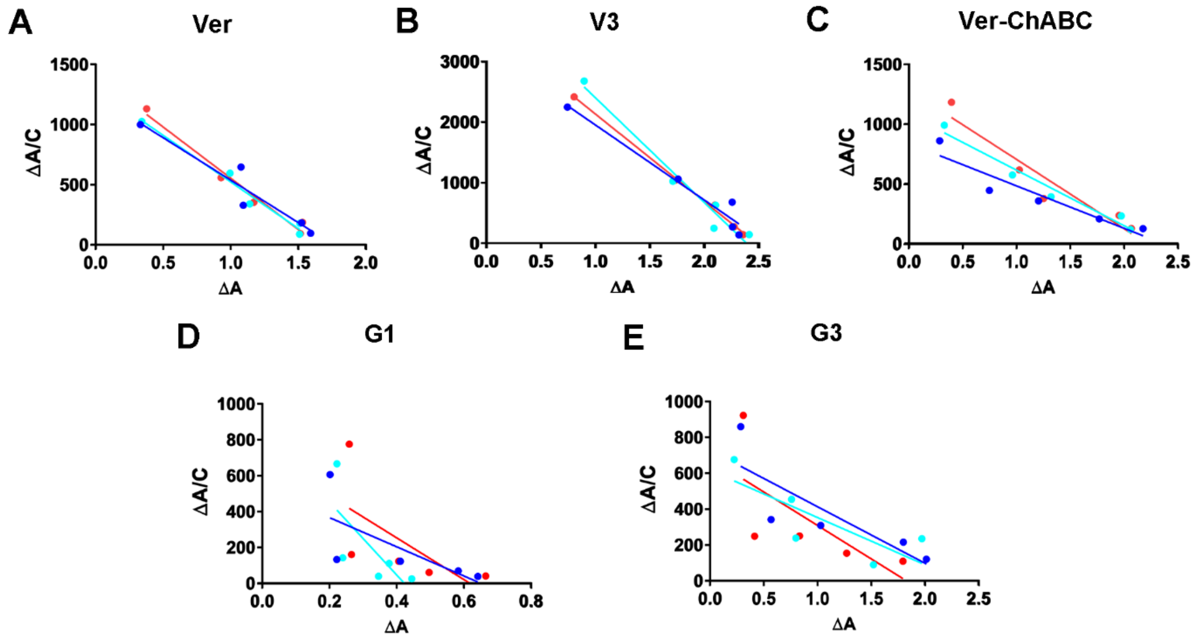


**Figure S3.** Scatchard analysis of solid phase binding data shown in Figure 2B & C. **A-E)** The Scatchard plot of the binding data from collagen-versican (A), collagen-V3 isoform (B), collagen-versican digested with chondroitinase ABC (C), collagen-G1 domain (D) and collagen-G3 domain (E). Three independent experiments were carried out for each.

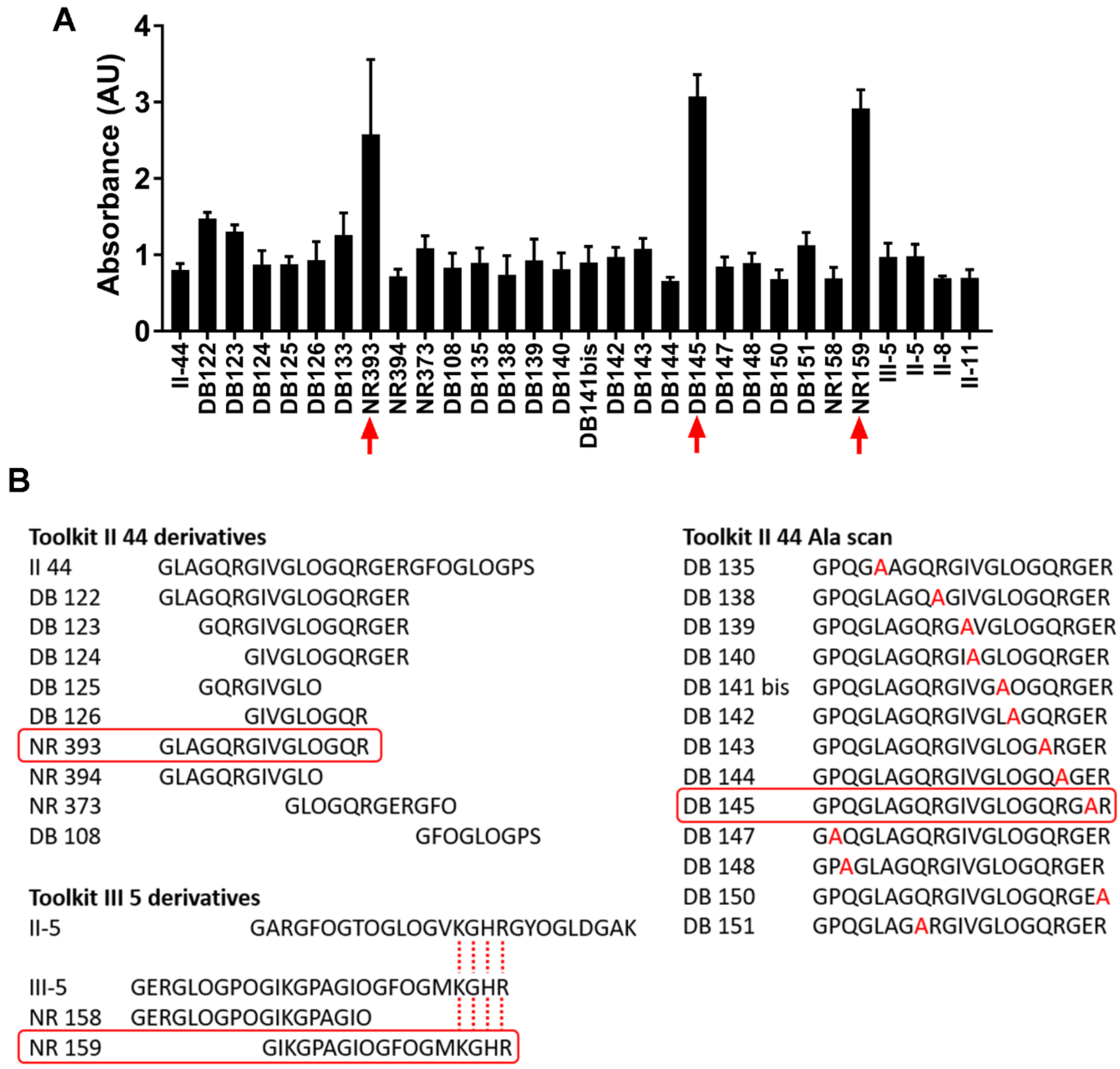


**Figure S4.** Versican G3 domain binds specific peptides with different motifs than the versican V3 isoform binds. **A)** Binding between variants of peptides II-44 and III-5 and 10 µg/ml recombinant G3 was tested using a solid-phase binding assay. **B)** The sequences of peptide II-4 and III-5 variants and the sequences of peptide II-4 modified with alanine (Ala/A, red). The high binding peptides are shown surrounded by a red box and the binding motifs (KGHR) found for peptide II-5, III-5 and III-5 variant are aligned with red dashed lines. Two independent experiments with three technical repeats each were carried out; data represent mean ± SD.

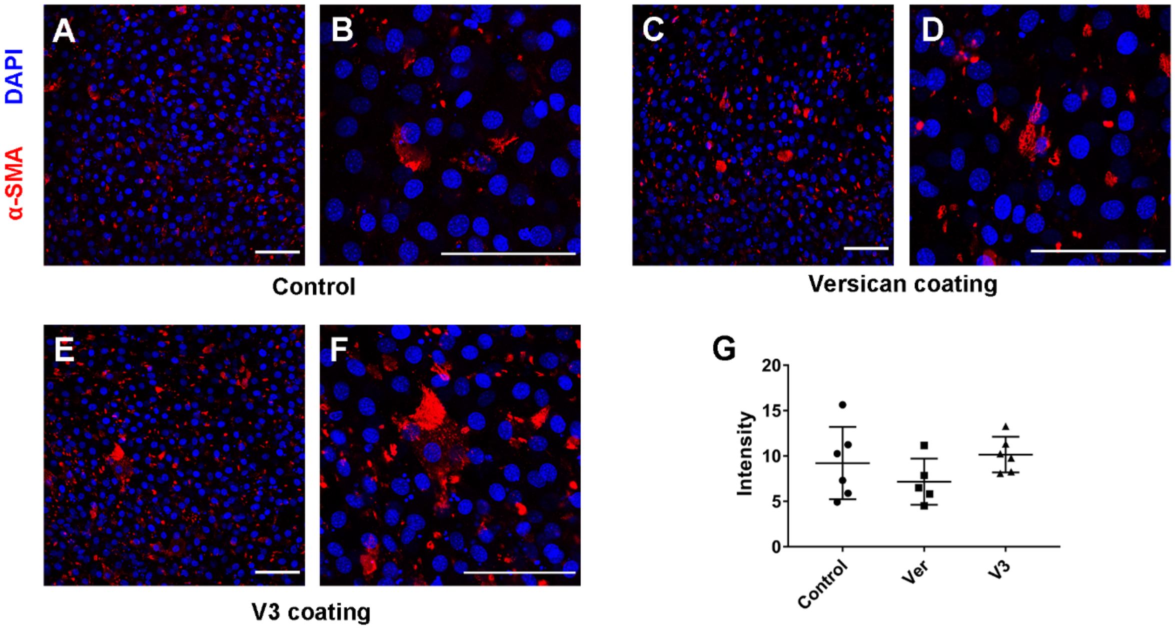


**Figure S5.** 3T3 fibroblasts on fibroblast-derived matrices are not activated due to the presence of versican or V3 isoform. **A-F)** Representative confocal imaging fibroblast-derived matrices immunostained for α-smooth muscle actin (α-SMA): vitronectin coating on plate as a control (A, B); versican coating (C, D); V3 coating (E, F). Scale bar = 200 μm. **G)** Quantification of α-SMA staining. One technical repeat from one individual experiment was used for α-SMA, and data points in (G) represent the intensity of α-SMA from images taken from each condition (6 images for control; 5 images for versican coating; 6 images for V3 coating).

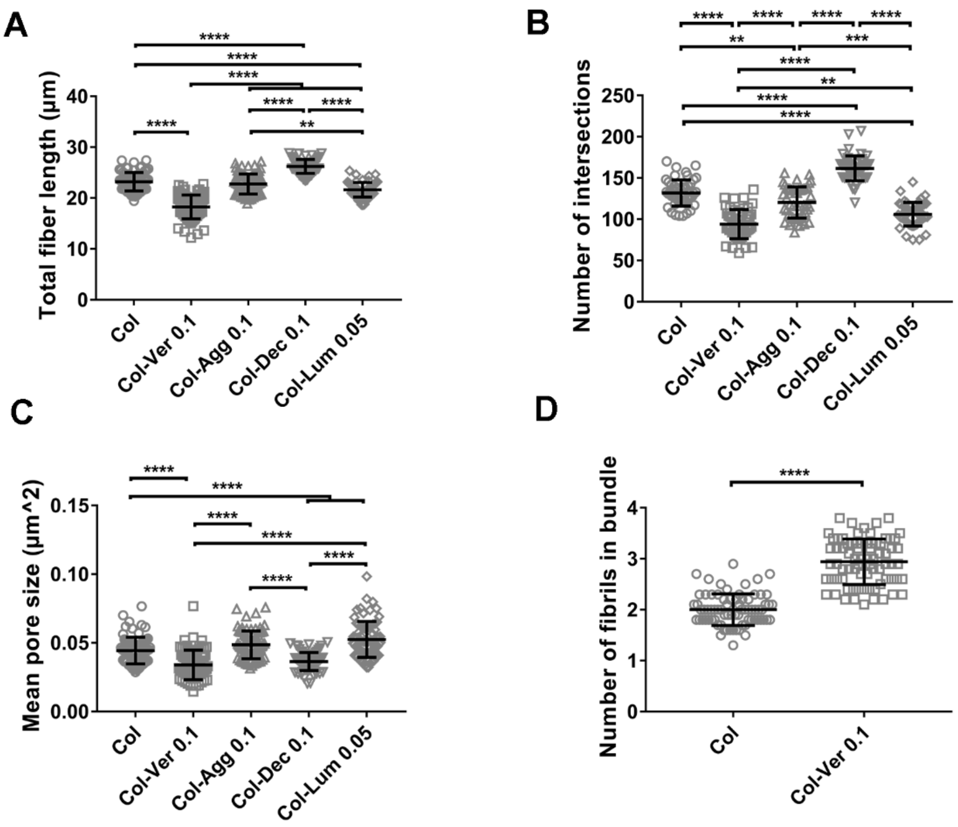


**Figure S6.** Versican and collagen form a loosely connected network with large fiber bundles and small pores. **A-C)** The total fiber length, number of intersections and mean pore size of collagen gels with different proteoglycans were quantified by DiameterJ using previously published SEM images [1]. Versican (Ver), aggrecan (Agg) and decorin (Dec) were added at 0.1 mg/ml to 1.5 mg/ml collagen (Col)and lumican (Lum) was added at 0.05 mg/ml to 1.5 mg/ml collagen (Col). ( **D)** The number of individual fibrils in bundles was counted manually from collagen and collagen-versican SEM images as we observed that versican increased collagen fibrils fusion into bundles.

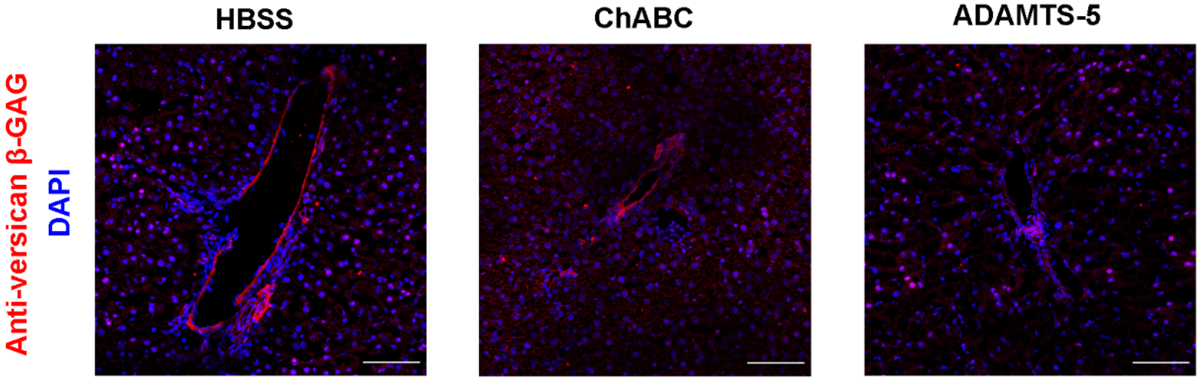


**Figure S7.** ADAMTS-5 perfusion of liver tissues effectively cleaves versican core protein. **A-C)** Representative confocal images of versican β-GAG-stained tissue in HBSS (A), ADAMTS-5 (B) and ChABC (C) perfused livers. L indicates lumen of a portal tract or vessel. Scale bar = 200 μm. Anti-versican staining (red), DAPI (blue). Anti-versican βGAG antibody targets amino acids 1360-1439 in the full-length protein, covering the ADAMTS-5 cleavage site, and therefore only stains intact versican.

**Table S1.** The amino acid sequences and molecular weight of 56 Collagen Toolkit II peptides. O represents hydroxyproline.

| Peptide | Sequence | MW |
|---|---|---|
| TK-II-1 | GPC-$(GPP)_5$-GPMGPMGPRGPOGPAGAOGPQGFQGNO-$(GPP)_5$-GPC-$NH_2$ | 5558 |
| TK-II-2 | GPC-$(GPP)_5$-GPQGFQGNOGEOGEOGVSGPMGPRGPO-$(GPP)_5$-GPC-$NH_2$ | 5648 |
| TK-II-3 | GPC-$(GPP)_5$-GPMGPRGPOGPOGKOGDDGEAGKOGKA-$(GPP)_5$-GPC-$NH_2$ | 5572 |
| TK-II-4 | GPC-$(GPP)_5$-GEAGKOGKAGERGPOGPQGARGFOGTO-$(GPP)_5$-GPC-$NH_2$ | 5621 |
| TK-II-5 | GPC-$(GPP)_5$-GARGFOGTOGLOGVKGHRGYOGLDGAK-$(GPP)_5$-GPC-$NH_2$ | 5710 |
| TK-II-6 | GPC-$(GPP)_5$-GYOGLDGAKGEAGAOGVKGESGSOGEN-$(GPP)_5$-GPC-$NH_2$ | 5533 |
| TK-II-7 | GPC-$(GPP)_5$-GESGSOGENGSOGPMGPRGLOGERGRT-$(GPP)_5$-GPC-$NH_2$ | 5668 |
| TK-II-8 | GPC-$(GPP)_5$-GLOGERGRTGPAGAAGARGNDGQOGPA-$(GPP)_5$-GPC-$NH_2$ | 5503 |
| TK-II-9 | GPC-$(GPP)_5$-GNDGQOGPAGPOGPVGPAGGOGFOGAO-$(GPP)_5$-GPC-$NH_2$ | 5385 |
| TK-II-10 | GPC-$(GPP)_5$-GGOGFOGAOGAKGEAGPTGARGPEGAQ-$(GPP)_5$-GPC-$NH_2$ | 5423 |
| TK-II-11 | GPC-$(GPP)_5$-GARGPEGAQGPRGEOGTOGSOGPAGAS-$(GPP)_5$-GPC-$NH_2$ | 5447 |
| TK-II-12 | GPC-$(GPP)_5$-GSOGPAGASGNOGTDGIOGAKGSAGAO-$(GPP)_5$-GPC-$NH_2$ | 5295 |
| TK-II-13 | GPC-$(GPP)_5$-GAKGSAGAOGIAGAOGFOGPRGPOGPQ-$(GPP)_5$-GPC-$NH_2$ | 5417 |
| TK-II-14 | GPC-$(GPP)_5$-GPRGPOGPQGATGPLGPKGQTGEOGIA-$(GPP)_5$-GPC-$NH_2$ | 5510 |
| TK-II-15 | GPC-$(GPP)_5$-GQTGEOGIAGFKGEQGPKGEOGPAGPQ-$(GPP)_5$-GPC-$NH_2$ | 5607 |
| TK-II-16 | GPC-$(GPP)_5$-GEOGPAGPQGAOGPAGEEGKRGARGEO-$(GPP)_5$-GPC-$NH_2$ | 5558 |
| TK-II-17 | GPC-$(GPP)_5$-GKRGARGEOGGVGPIGPOGERGAOGNR-$(GPP)_5$-GPC-$NH_2$ | 5628 |
| TK-II-18 | GPC-$(GPP)_5$-GERGAOGNRGFOGQDGLAGPKGAOGER-$(GPP)_5$-GPC-$NH_2$ | 5680 |
| TK-II-19 | GPC-$(GPP)_5$-GPKGAOGERGPSGLAGPKGANGDOGRO-$(GPP)_5$-GPC-$NH_2$ | 5529 |
| TK-II-20 | GPC-$(GPP)_5$-GANGDOGROGEOGLOGARGLTGROGDA-$(GPP)_5$-GPC-$NH_2$ | 5606 |
| TK-II-21 | GPC-$(GPP)_5$-GLTGROGDAGPQGKVGPSGAOGEDGRO-$(GPP)_5$-GPC-$NH_2$ | 5562 |
| TK-II-22 | GPC-$(GPP)_5$-GAOGEDGROGPOGPQGARGQOGVMGFO-$(GPP)_5$-GPC-$NH_2$ | 5650 |
| TK-II-23 | GPC-$(GPP)_5$-GQOGVMGFOGPKGANGEOGKAGEKGLO-$(GPP)_5$-GPC-$NH_2$ | 5625 |
| TK-II-24 | GPC-$(GPP)_5$-GKAGEKGLOGAOGLRGLOGKDGETGAA-$(GPP)_5$-GPC-$NH_2$ | 5536 |
| TK-II-25 | GPC-$(GPP)_5$-GKDGETGAAGPOGPAGPAGERGEQGAO-$(GPP)_5$-GPC-$NH_2$ | 5447 |
| TK-II-26 | GPC-$(GPP)_5$-GERGEQGAOGPSGFQGLOGPOGPOGEG-$(GPP)_5$-GPC-$NH_2$ | 5577 |
| TK-II-27 | GPC-$(GPP)_5$-GPOGPOGEGGKOGDQGVOGEAGAOGLV-$(GPP)_5$-GPC-$NH_2$ | 5458 |
| TK-II-28 | GPC-$(GPP)_5$-GEAGAOGLVGPRGERGFOGERGSOGAQ-$(GPP)_5$-GPC-$NH_2$ | 5638 |
| TK-II-29 | GPC-$(GPP)_5$-GERGSOGAQGLQGPRGLOGTOGTDGPK-$(GPP)_5$-GPC-$NH_2$ | 5917 |
| TK-II-30 | GPC-$(GPP)_5$-GTOGTDGPKGASGPAGPOGAQGPOGLQ-$(GPP)_5$-GPC-$NH_2$ | 5401 |
| TK-II-31 | GPC-$(GPP)_5$-GAQGPOGLQGMOGERGAAGIAGPKGDR-$(GPP)_5$-GPC-$NH_2$ | 5561 |
| TK-II-32 | GPC-$(GPP)_5$-GIAGPKGDRGDVGEKGPEGAOGKDGGR-$(GPP)_5$-GPC-$NH_2$ | 5525 |
| TK-II-33 | GPC-$(GPP)_5$-GAOGKDGGRGLTGPIGPOGPAGANGEK-$(GPP)_5$-GPC-$NH_2$ | 5444 |
| TK-II-34 | GPC-$(GPP)_5$-GPAGANGEKGEVGPOGPAGSAGARGAO-$(GPP)_5$-GPC-$NH_2$ | 5344 |
| TK-II-35 | GPC-$(GPP)_5$-GSAGARGAOGERGETGPOGPAGFAGPO-$(GPP)_5$-GPC-$NH_2$ | 5450 |
| TK-II-36 | GPC-$(GPP)_5$-GPAGFAGPOGADGQOGAKGEQGEAGQK-$(GPP)_5$-GPC-$NH_2$ | 5495 |
| TK-II-37 | GPC-$(GPP)_5$-GEQGEAGQKGDAGAOGPQGPSGAOGPQ-$(GPP)_5$-GPC-$NH_2$ | 5475 |
| TK-II-38 | GPC-$(GPP)_5$-GPSGAOGPQGPTGVTGPKGARGAQGPO-$(GPP)_5$-GPC-$NH_2$ | 5412 |
| TK-II-39 | GPC-$(GPP)_5$-GARGAQGPOGATGFOGAAGRVGPOGSN-$(GPP)_5$-GPC-$NH_2$ | 5436 |
| TK-II-40 | GPC-$(GPP)_5$-GRVGPOGSNGNOGPOGPOGPSGKDGPK-$(GPP)_5$-GPC-$NH_2$ | 5525 |
| TK-II-41 | GPC-$(GPP)_5$-GPSGKDGPKGARGDSGPOGRAGEOGLQ-$(GPP)_5$-GPC-$NH_2$ | 5561 |
| TK-II-42 | GPC-$(GPP)_5$-GRAGEOGLQGPAGPOGEKGEOGDDGPS-$(GPP)_5$-GPC-$NH_2$ | 5561 |
| TK-II-43 | GPC-$(GPP)_5$-GEOGDDGPSGAEGPOGPQGLAGQRGIV-$(GPP)_5$-GPC-$NH_2$ | 5531 |
| TK-II-44 | GPC-$(GPP)_5$-GLAGQRGIVGLOGQRGERGFOGLOGPS-$(GPP)_5$-GPC-$NH_2$ | 5705 |
| TK-II-45 | GPC-$(GPP)_5$-GFOGLOGPSGEOGKQGAOGASGDRGPO-$(GPP)_5$-GPC-$NH_2$ | 5551 |
| TK-II-46 | GPC-$(GPP)_5$-GASGDRGPOGPVGPOGLTGPAGEOGRE-$(GPP)_5$-GPC-$NH_2$ | 5514 |
| TK-II-47 | GPC-$(GPP)_5$-GPAGEOGREGSOGADGPOGRDGAAGVK-$(GPP)_5$-GPC-$NH_2$ | 5491 |
| TK-II-48 | GPC-$(GPP)_5$-GRDGAAGVKGDRGETGAVGAOGAOGPO-$(GPP)_5$-GPC-$NH_2$ | 5449 |
| TK-II-49 | GPC-$(GPP)_5$-GAOGAOGPOGSOGPAGPTGKQGDRGEA-$(GPP)_5$-GPC-$NH_2$ | 5431 |
| TK-II-50 | GPC-$(GPP)_5$-GKQGDRGEAGAQGPMGPSGPAGARGIQ-$(GPP)_5$-GPC-$NH_2$ | 5534 |

| | | |
|---|---|---|
| TK-II-51 | GPC-$(GPP)_5$-GPAGARGIQGPQGPRGDKGEAGEOGER-$(GPP)_5$-GPC-$NH_2$ | 5644 |
| TK-II-52 | GPC-$(GPP)_5$-GEAGEOGERGLKGHRGFTGLQGLOGPO-$(GPP)_5$-GPC-$NH_2$ | 5746 |
| TK-II-53 | GPC-$(GPP)_5$-GLQGLOGPOGPSGDQGASGPAGPSGPR-$(GPP)_5$-GPC-$NH_2$ | 5427 |
| TK-II-54 | GPC-$(GPP)_5$-GPAGPSGPRGPOGPVGPSGKDGANGIO-$(GPP)_5$-GPC-$NH_2$ | 5409 |
| TK-II-55 | GPC-$(GPP)_5$-GKDGANGIOGPIGPOGPRGRSGETGPA-$(GPP)_5$-GPC-$NH_2$ | 5528 |
| TK-II-56 | GPC-$(GPP)_5$-GPRGRSGETGPAGPOGNOGPOGPOGPO-$(GPP)_5$-GPC-$NH_2$ | 5521 |

**Table S2.** Statistical significance of differences in fibril orientation between conditions. Data from Figure 4E were analyzed using two-way ANOVA with repeated measurements. *P<0.05, **P<0.01, ***P<0.001 and ****P<0.0001.

| Data #1 | Data #2 | Angle range | Significance |
|---|---|---|---|
| Control | Ver | -40° to -30° | ** |
| Control | Ver | -30° to -20° | **** |
| Control | Ver | -20° to -10° | **** |
| Control | Ver | -10° to 0° | **** |
| Control | Ver | 0° to 10° | **** |
| Control | Ver | 10° to 20° | *** |
| Control | Ver | 20° to 30° | *** |
| Control | Ver | 30° to 40° | ** |
| Control | V3 | -40° to -30° | * |
| Control | V3 | -30° to -20° | * |
| Control | V3 | -20° to -10° | * |
| Control | V3 | 20° to 30° | ** |
| Control | V3 | 30° to 40° | ** |
| Control | V3 | 40° to 50° | ** |
| Control | V3 | 50° to 60° | * |
| Ver | V3 | -30° to -20° | * |
| Ver | V3 | -20° to -10° | *** |
| Ver | V3 | -10° to 0° | *** |
| Ver | V3 | 0° to 10° | ** |

**Table S3.** Statistical significance of differences in G' for different collagen co-gels subjected to a strain sweep test. Data were analyzed using two-way ANOVA with repeated measurements. *P<0.05, **P<0.01, ***P<0.001 and ****P<0.0001.

| Data #1 | Data #2 | Shear strain (%) | Significance |
|---|---|---|---|
| Col | Col-Ver | 7.9433 | ** |
| Col | Col-Ver | 10 | **** |
| Col | Col-Ver | 12.5893 | **** |
| Col | Col-Ver | 15.8489 | **** |
| Col | Col-Ver | 19.9625 | **** |
| Col | Col-Ver | 25.1189 | **** |
| Col | Col-V3 | 1 | * |
| Col | Col-V3 | 10 | * |
| Col | Col-V3 | 12.5893 | **** |
| Col | Col-V3 | 15.8489 | **** |
| Col | Col-V3 | 19.9625 | **** |
| Col | Col-V3 | 25.1189 | **** |
| Col | Col-Agg | 15.8489 | ** |
| Col | Col-Agg | 19.9625 | **** |
| Col | Col-Agg | 25.1189 | **** |
| Col | Col-Dec | 19.9625 | **** |
| Col | Col-Dec | 25.1189 | **** |
| Col-Ver | Col-V3 | 7.9433 | * |
| Col-Ver | Col-Agg | 7.9433 | * |
| Col-Ver | Col-Agg | 10 | **** |
| Col-Ver | Col-Agg | 12.5893 | **** |
| Col-Ver | Col-Agg | 15.8489 | **** |
| Col-Ver | Col-Agg | 19.9625 | **** |
| Col-Ver | Col-Agg | 25.1189 | * |
| Col-Ver | Col-Dec | 7.9433 | ** |
| Col-Ver | Col-Dec | 10 | **** |
| Col-Ver | Col-Dec | 12.5893 | **** |
| Col-Ver | Col-Dec | 15.8489 | **** |
| Col-V3 | Col-Dec | 10 | ** |
| Col-V3 | Col-Dec | 12.5893 | **** |
| Col-V3 | Col-Dec | 15.8489 | **** |
| Col-V3 | Col-Agg | 12.5893 | **** |
| Col-V3 | Col-Agg | 15.8489 | **** |
| Col-V3 | Col-Agg | 19.9625 | ** |
| Col-Agg | Col-Dec | 15.8489 | ** |
